# Cdc73 majorly regulates apoptosis-inducing factor (*AIF1*) in *Saccharomyces cerevisiae* via the H3K36 methylation

**DOI:** 10.1101/2023.11.06.565826

**Authors:** Nitu Saha, Santoshi Acharjee, Raghuvir Singh Tomar

## Abstract

*AIF1* overexpression is intimately linked to the sensitivity of the yeast cells towards hydrogen peroxide or acetic acid. Therefore, studying the mechanism of its regulation in the cell would provide a significant understanding of the factors ultimately guiding yeast apoptosis. In this report, we establish the time-dependent induction of *AIF1* in hydrogen peroxide stress. Additionally, the *AIF1* expression in hydrogen peroxide is mediated by two transcription factors, Yap5 (DNA binding) and Cdc73 (non-DNA binding). Furthermore, substituting the H3K36 residue with another significantly abrogates the *AIF1* expression. However, substituting H3K4 or H3K79 with A does not affect the *AIF1* expression level under hydrogen peroxide stress. Altogether, the significant reduction of *AIF1* expression in *cdc73*Δ cells plausibly reflects the reduced H3K36me3 modification and is independent of the H3K4me3 modification.

## Introduction

Regulated cell death (RCD) in yeast cells due to apoptotic insults is executed by distinct mechanisms of different genes such as *NMA111* [1], *YCA1* [2], *NUC1* [3], *BXI1* [4], and *AIF1* [5]. Initial studies of these genes have given insights into the role mediated by these apoptotic proteins. However, the factors regulating the gene expression levels of these apoptotic genes have yet to be studied. Studies by Wissing et al. [5] have thoroughly investigated the role of DNA fragmentation by Aif1p after translocation from the mitochondria to the nucleus in yeast apoptosis. Further observations revealed that hydrogen peroxide treatment accumulates the Aif1p levels. However, the precise reason behind the accumulation of Aif1 still needs to be explored.

The methylation of lysine residue in histones controls the transcription process. Furthermore, the modifications occur primarily during the initiation or elongation phases of transcription by RNA polymerase. H3K36 methylation and active transcriptional elongation process are coupled with the gradient of H3K36 methylation increasing towards the 3’ end of the gene. Set2, a specific histone methylase for the H3K36 residue, preferentially associates with elongating RNA polymerase II form.

Yap5, a transcription factor, regulates the expression of genes like *CCC1* and *GRX4,* which maintain iron homeostasis in replete conditions. The regulation is primarily through binding the consensus sequence called YREs (Yap1 recognition element) at the promoter regions of the gene [6]. Deletion of *yap5* results in a resistant phenotype against sorbate or benzoate primarily through the induction of Pdr12p, a plasma membrane transporter protein [7].

Cdc73, a non-DNA binding transcription factor, is a Paf1 (RNA Polymerase II Associated Factor) complex component. The other components are Ctr9, Rtf1, Paf1 and Leo1 [8, 9]. The Cdc73 subunit is highly conserved across different species, and its deletion is detrimental to the transcription elongation efficiency of the complex [10, 11]. *cdc73*Δ reduces the interaction between Paf1C and RNA polymerase II and reduces the localisation of Paf1C at the transcribed genes [12]. Paf1C also regulates multiple histone modifications, such as methylation of H3K4, H3K79 and H3K36 [13–15]. Particularly, the Cdc73 subunit maintains the H3K36me3 signatures on the coding regions of the genes which are active and the H3K4me3 but not H3K4me2 or H3K79me2/H3K79me3 [16]. Together, the studies show that Cdc73 regulates the transcription of many genes in the yeast by impacting the interaction between the Paf1C and RNA polymerase II and the histone modifications [16].

The histone methylation network in *Saccharomyces cerevisiae* is vast and caters to regulating multiple physiologically crucial pathways. Different methylation marks are associated with different patterns of gene expression. Of various methylation marks, the H3K4, H3K36 and H3K79 are associated with euchromatin, the transcriptionally active region of the chromatin. H3K4 residue can methylate at three levels to form H3K4me, H3K4me2 and H3K4me3 [17]. Monomethylated H3K4 is enriched at the 3’-end of the transcriptional genes [18–20], whereas the dimethylated versions are enriched in the coding regions of the transcriptionally active or poised areas [21]. The trimethylated version is enriched in the promoter regions of regulated genes and at the 5’ end of the genes undergoing active transcription [21]. Since the marks are so intimately linked with transcription, the yeast cells with defective methylation of H3K4 exhibit multiple problems of decreased chronological ageing [22], faulty DNA repair systems [23], and sensitivity to many antifungal drugs [24].

Like H3K4 residue, yeast cells co-transcriptionally exhibit H3K36 mono/di/trimethylation marks. H3K36 monomethylation is concentrated at the 5’-end of the genes [25] showing transcription, whereas di- and trimethylation are concentrated towards the 3’-ends. Of the different methylations at the H3K36 residue, the trimethylated version exhibits a high correlation with actively transcribed regions and ensures the permissive transcription is maintained by recruiting reader proteins [26, 27]. Thus, the methylation of H3K36 is crucial for DNA damage repair and transcriptional regulation [28–30].

Unlike the other two methylation sites in the histone tail region, the H3K79 residue is present at the globular core of the nucleosome [31]. The H3K79 residue exhibits methylation at three levels viz., H3K79me1/2/3, of which the trimethylated version is the most prevalent form [32]. The H3K79 methylation, unlike the H3K4 or H3K36, does not show enrichment on one specific region but is dispersed uniformly across the actively transcribed gene coding region [25, 33]. Methylated H3K79 residue interacts with the Sir proteins to establish the heterochromatin regions and maintenance of silent telomeric regions [34]. The modification also interacts with Rad9p, which inhibits the single-stranded DNA (ssDNA) production at the double-strand break (DSB) site, thus playing a crucial role in DNA damage repair [35, 36].

Previously, it was shown that the Aif1p increased upon hydrogen peroxide treatment in yeast cells. However, the reason behind the increase remains unknown [5]. In this communication, we, for the first time, show that treatment with hydrogen peroxide selectively increases the *AIF1* gene expression level but not other apoptotic genes. Furthermore, we show that the increase depends on the transcription factor Yap5 and Cdc73, a component of the Polymerase-Associated Factor 1 (Paf1) complex (Paf1C). *yap5*Δ or *cdc73*Δ reduces the *AIF1* gene expression upon hydrogen peroxide treatment. Furthermore, the histone methylation at the K36 residue is critical for maintaining basal and hydrogen peroxide-induced *AIF1* gene induction, as the H3K36A mutant shows significantly lower levels of the gene. Thus, our study demonstrates the crucial role of histone modifications in the maintenance of *AIF1* gene expression.

## Results

### *AIF1* gene expression selectively increases during hydrogen peroxide stress

Immunoblot studies by Wissing et al. revealed that hydrogen peroxide treatment increases the Aif1 levels [5]. We probed whether an increase in the Aif1p level reflects the increased mRNA levels of the gene. To this end, we treated the wild-type (WT) cells with different concentrations of hydrogen peroxide at 0.8 and 1 mM and harvested cells at different time points of 15’, 30’ and 45’. We found that compared to the untreated (UT) condition, at 15’, the *AIF1* gene expression levels increased by 7.5±1.6-fold and 3.9±1.1-fold; at 30’, the increase was about 12.7±1.7-fold and 7.3±3.5-fold and at 45’, the increase was about 10.1±2 or 8.6±2.8-fold, respectively for 0.8 mM and 1 mM concentration of H_2_O_2_ treatment (Fig. 1A). Next, we checked the expression levels of other apoptotic genes viz. *YCA1*, *NUC1*, *NMA111* and *BXI1* upon exposure to hydrogen peroxide under the same conditions. We found that compared to UT the levels of *YCA1* (0.95±0.04 and 0.98±0.06, 1.23±0.12 and 1.03±0.23, 0.91±0.40 and 0.77±0.33 for 15’, 30’ and 45’, respectively), *NUC1* (0.75±0.08 and 0.90±0.15, 0.94±0.11 and 0.80±0.21, 0.90±0.39 and 0.64±0.04 for 15’, 30’ and 45’, respectively), *NMA111* (0.74±0.05 and 0.72±0.07, 1.18±0.15 and 0.85±0.11, 1.05±0.52 and 0.69±0.28 for 15’, 30’ and 45’, respectively) did not exhibit substantial change (Fig. 1B, C and D). The *BXI1* gene expression level (2.35±0.64 and 1.51±0.30, 3.35±1.21 and 3.16±1.90, 3.8±0.37 and 3.74±1.50 for 15’, 30’ and 45’, respectively) increases significantly at 45’ with 0.8 mM H_2_O_2_ treatment (Fig. 1E). Together, the results indicate that *AIF1* gene expression levels increase significantly and selectively upon hydrogen peroxide stress.

**Figure 1.**
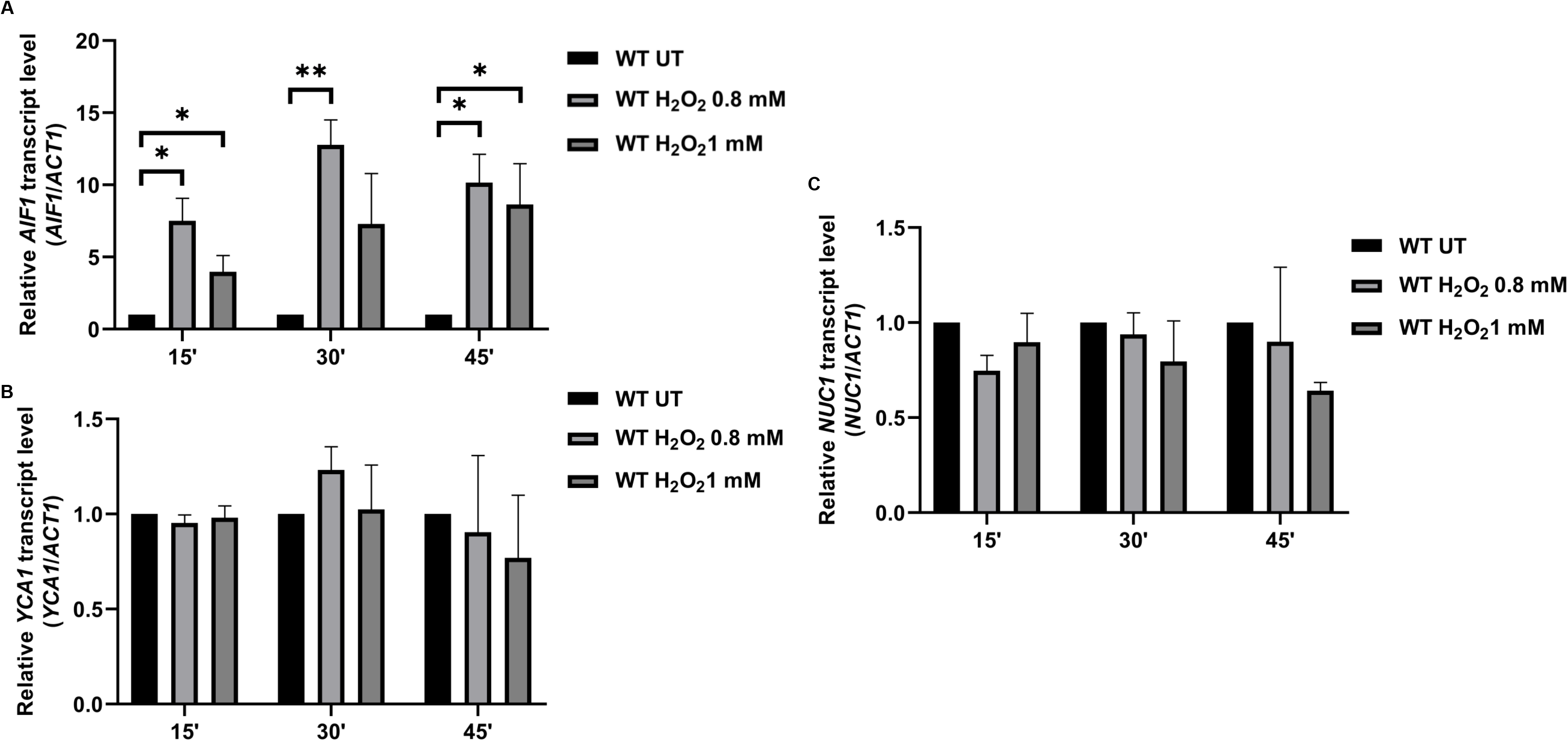
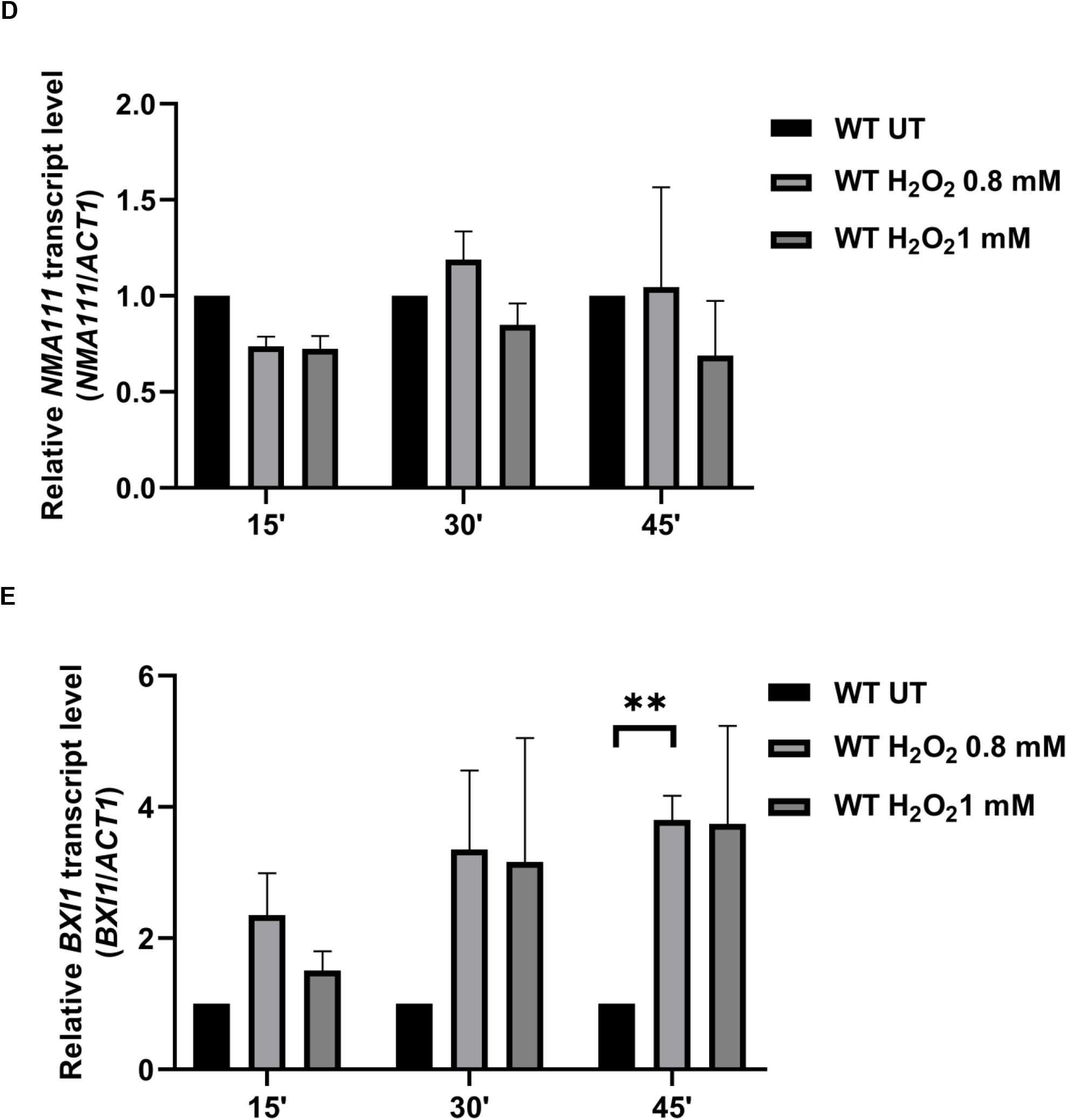
Treatment with hydrogen peroxide selectively induces the expression of *AIF1* gene levels. **A,** *AIF1* or **B,** *YCA1* **C,** *NUC1* **D,** *NMA111* or **E,** *BXI1* mRNA level was examined in the wild-type cells (H3 WT) under untreated (UT) or H_2_O_2_ treated conditions (0.8 mM, 1 mM) for 15’, 30’ or 45’. The relative expression with respect to the *ACT1* gene (control) is shown. Statistical analysis was performed using Student’s t-test (GraphPad Prism 8 GraphPad Software, Inc). Values shown are the mean ± SD of three biological replicates (n = 3). ∗∗p (0.001 to 0.01), ∗p (0.01 to 0.05)

### *AIF1* gene expression plays a crucial role in determining the cell survival under hydrogen peroxide stress

*AIF1* gene expression level is regulated by two transcription factors, a DNA binding Yap5p and a non-DNA binding Cdc73p (Fig. 2A) [37, 38]. However, the role of these transcription factors in the regulation of the *AIF1* gene upon hydrogen peroxide treatment is unknown. Thus, we created deletions of Yap5 and/or Cdc73 in the WT background (Fig. S1). Since *AIF1* gene deletion is resistant to hydrogen peroxide stress, an apoptotic agent, we first tested if the strains deleted for the *AIF1* regulators showed differential resistance to H_2_O_2_. To this end, the strains were spotted in increasing concentrations of H_2_O_2_ (0.8, 1, 1.25 and 1.5 mM). We found that *yap5*Δ or *aif1*Δ grew better than the WT cells at 1.25 mM concentration, whereas *cdc73*Δ sensitivity at 1.5 mM concentration (Fig. 2B). We reconfirmed our observations by performing the growth curve kinetic studies. To this end, the cells from the primary culture were secondary inoculated at 0.2 OD in liquid culture under UT or H_2_O_2_-treated conditions and the growth was checked every 30 min. The growth curves revealed that at 3 mM concentration, the *aif1*Δ or *yap5*Δ cells grew better than WT (Fig. 2C, E) while *cdc73*Δ cells failed to grow. We repeated the growth assay for *cdc73*Δ cells at a lower concentration (2 mM) of hydrogen peroxide to find that the cells were indeed sensitive to H_2_O_2_ treatment compared to the WT cells. We also tested if the deleted cells behave similarly in the presence of another apoptotic agent, acetic acid, via spot test assay and growth curve assay. Fig. S2 A and B show that *aif1*Δ or *yap5*Δ cells grew better than WT cells in acetic acid at 80, 90 and 100 mM concentrations, whereas the *cdc73*Δ showed slower growth. Thus, the *yap5*Δ cells exhibit better growth in hydrogen peroxide and acetic acid, whereas the *cdc73*Δ cells do not exhibit the same phenotype. The sensitivity of *cdc73*Δ cells could reflect its role in regulating many gene expressions, which may be crucial for the survival of cells in the presence of multiple stresses. We also checked the survival of cells through chronological life span (CLS) assay and found no differential survival of cells during the 20-day span (Fig. S3). **Cdc73 majorly regulates the *AIF1* gene expression level under hydrogen peroxide stress.** Chromatin immunoprecipitation studies have revealed that Yap5 regulates the *AIF1* gene expression during heat stress [37]. Also, microarray analysis revealed that the cdc73 deletion changes the *AIF1* expression levels compared to the wild-type [38]. We argued that if these transcription factors regulate *AIF1* gene expression levels, it may regulate the expression pattern during the gene induction via hydrogen peroxide. To this end, we grew cells until the exponential phase and left them UT or treated them with 0.8 mM for 30’. Next, we extracted the RNA from the harvested cells and checked the *AIF1* gene expression from the cDNA obtained. As shown in Fig. 3A, compared to the WT (1.00±0.00) for UT and (15.16±0.41) for 0.8 mM H_2_O_2_ treatment, respectively, the *yap5*Δ cells show expression at (1.97±0.08 and 11.16±1.68 for UT and H_2_O_2_ treatment, respectively). Thus, the results indicate that the induction of *AIF1* gene expression upon H_2_O_2_ treatment is Yap5 dependent. Similarly, *cdc73*Δ cells significantly reduce the basal and H_2_O_2_-induced *AIF1* levels compared to the wild-type counterpart. As shown in Fig. 3B, compared to the WT (1.00±0.00) for UT and (19.97±2.57) for 0.8 mM H_2_O_2_ treatment, respectively, the *cdc73*Δ cells show expression at (0.37±0.10 and 5.49±1.86 for UT and H_2_O_2_ treatment, respectively). Thus, the results indicate that the induction of *AIF1* gene expression upon H_2_O_2_ treatment depends on Cdc73p.

**Figure 2.**
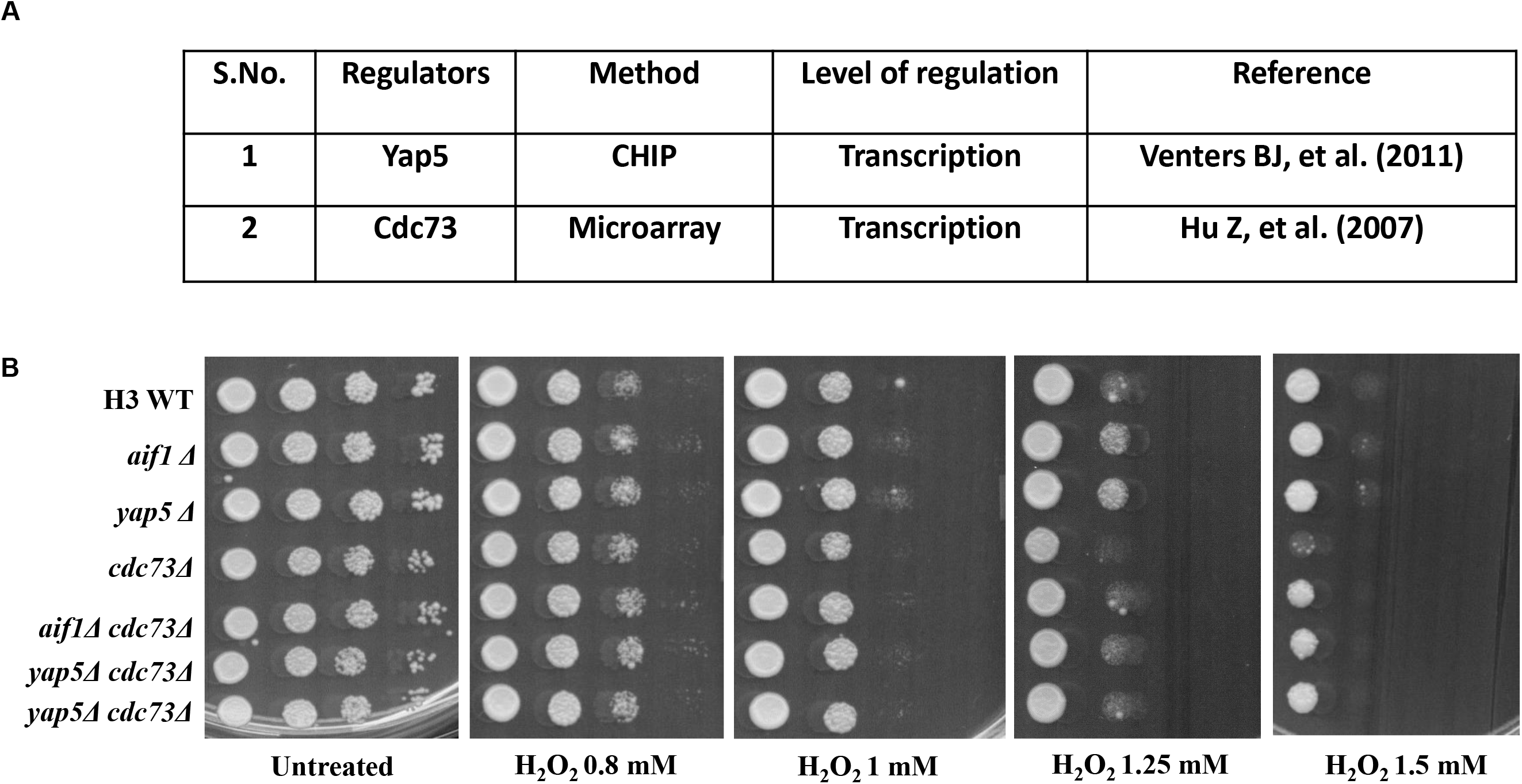
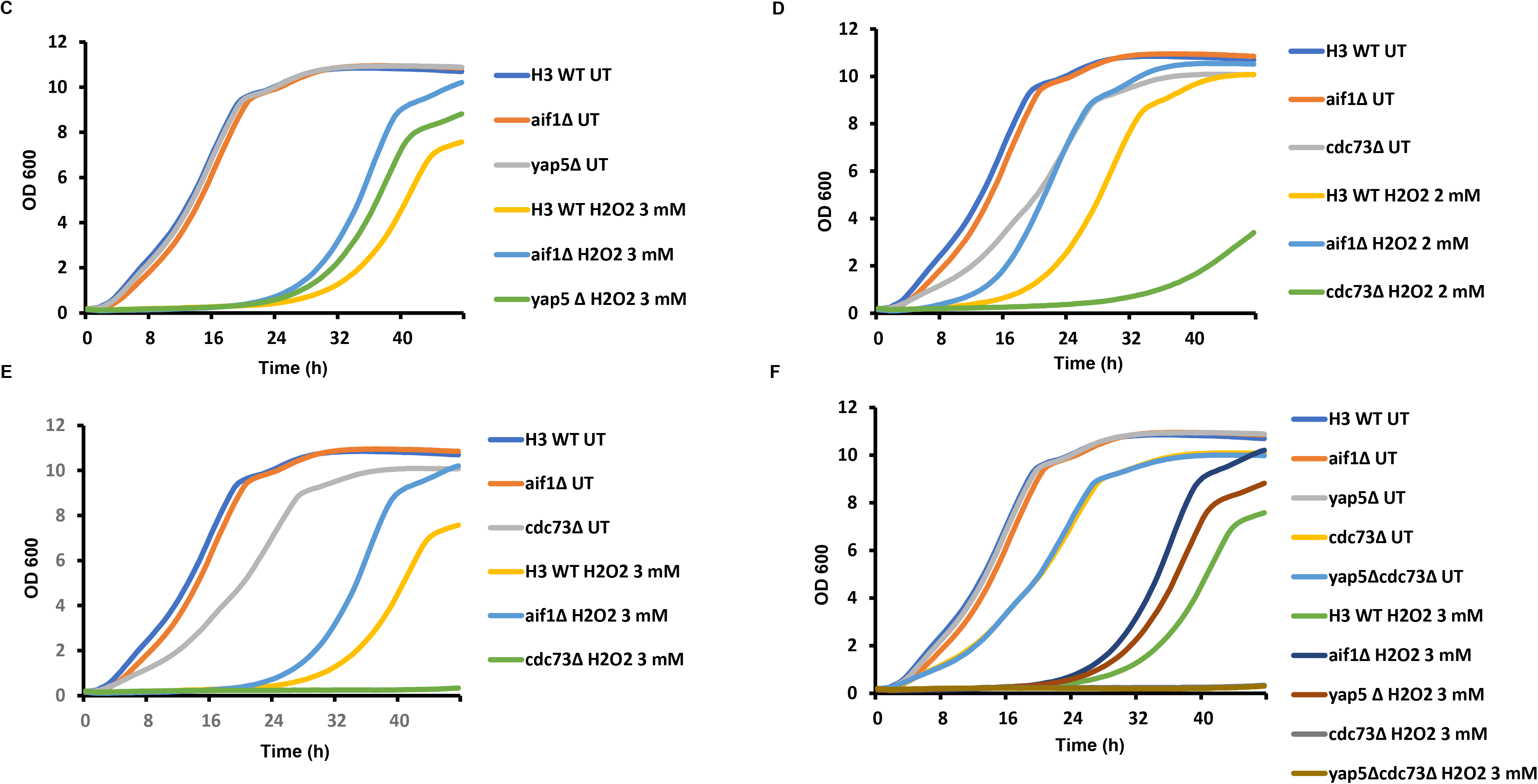
*AIF1* gene expression is crucial in determining cell survival under hydrogen peroxide stress. **A,** Summary of the known *AIF1* gene regulators. **B,** Spot test assay of the wild-type cells (H3 WT) and the indicated deletions of *aif1* or its regulators (at 10-fold dilution from left to right) in untreated (UT) or different H_2_O_2_ concentrations (0.8, 1, 1.25 and 1.5 mM). The untreated plate represents the SC + agar plate without any H_2_O_2_ and shows the growth of cells in the absence of stress. Cells were incubated at 30°C, and plates were scanned at 96 h. **C, D, E, F,** Growth of the H3 WT, and the indicated deletions in liquid culture under untreated (UT) or H_2_O_2_ treated (2 mM, 3 mM) conditions. The graphs reflect the absorbance values at 600 nm wavelength at every 30’ for the time indicated in the x-axis.

**Figure 3.**
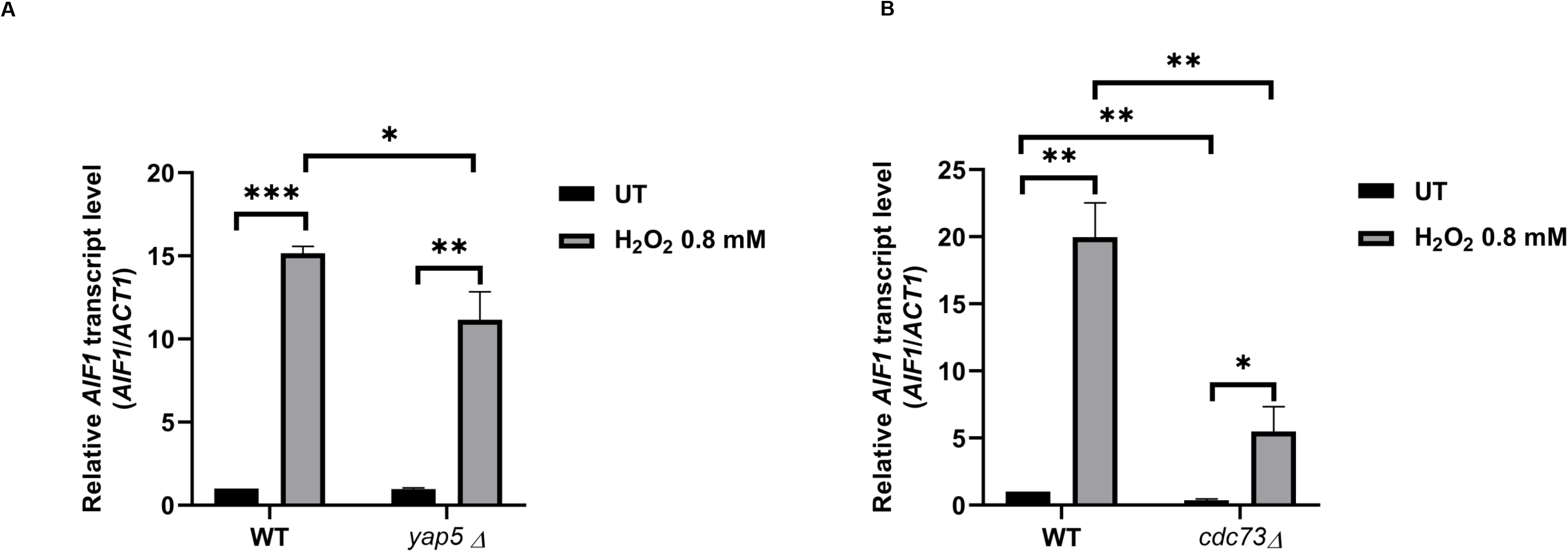
Cdc73 majorly regulates the *AIF1* gene expression level under hydrogen peroxide stress. **A,** *AIF1* mRNA level was examined in the wild-type cells (H3 WT) or *yap5*Δ under untreated (UT) or H_2_O_2_ treated conditions (0.8 mM) for 30’. **B,** *AIF1* mRNA level was examined in the wild-type cells (H3 WT) or *cdc73*Δ under untreated (UT) or H_2_O_2_ treated conditions (0.8 mM) for 30’. The relative expression with respect to the *ACT1* gene (control) is shown. Statistical analysis was performed using Student’s t-test (GraphPad Prism 8 GraphPad Software, Inc). The values are the mean ± SD of three replicates (n = 3). ∗∗∗p < 0.001, ∗∗p (0.001 to 0.01), ∗p (0.01 to 0.05).

### Global levels of H3K4me3 or H3K36me3 histone modifications remain unaltered under hydrogen peroxide stress

Since the *cdc73*Δ cells show low expression levels of the *AIF1* gene, we argued that hydrogen peroxide stress might be linked to the two histone modifications H3K4me3 and H3K36me3 levels which have been reported to be reduced in the *cdc73*Δ cells [16]. Thus, we checked the global levels of H3K4me3 and H3K36me3 after the H_2_O_2_ treatment. To this end, the WT or deleted strains were grown till the exponential phase and either kept untreated or treated with 0.8 mM H_2_O_2_ for 30’. The protein was extracted from the harvested cells and analysed using immunoblotting. As elucidated in Fig. 4, the WT or *yap5*Δ cells did not exhibit any change in the global H3K4me3 or H3K36me3 levels. As previously reported, the *cdc73*Δ cells show reduced levels of H3K4me3 and H3K36me3. However, no alteration was seen in H_2_O_2_ treatment. Furthermore, the *yap5*Δ *cdc73*Δ cells did not exhibit any change in H3K4me3 or H3K36me3 levels, indicating the effect on the reduced levels of these two modifications was only due to *cdc73*Δ in the double deletion. Thus, we conclude that the global modifications of H3K4me3 or H3K36me3 remain unaltered under the H_2_O_2_ stress.

**Figure 4.**
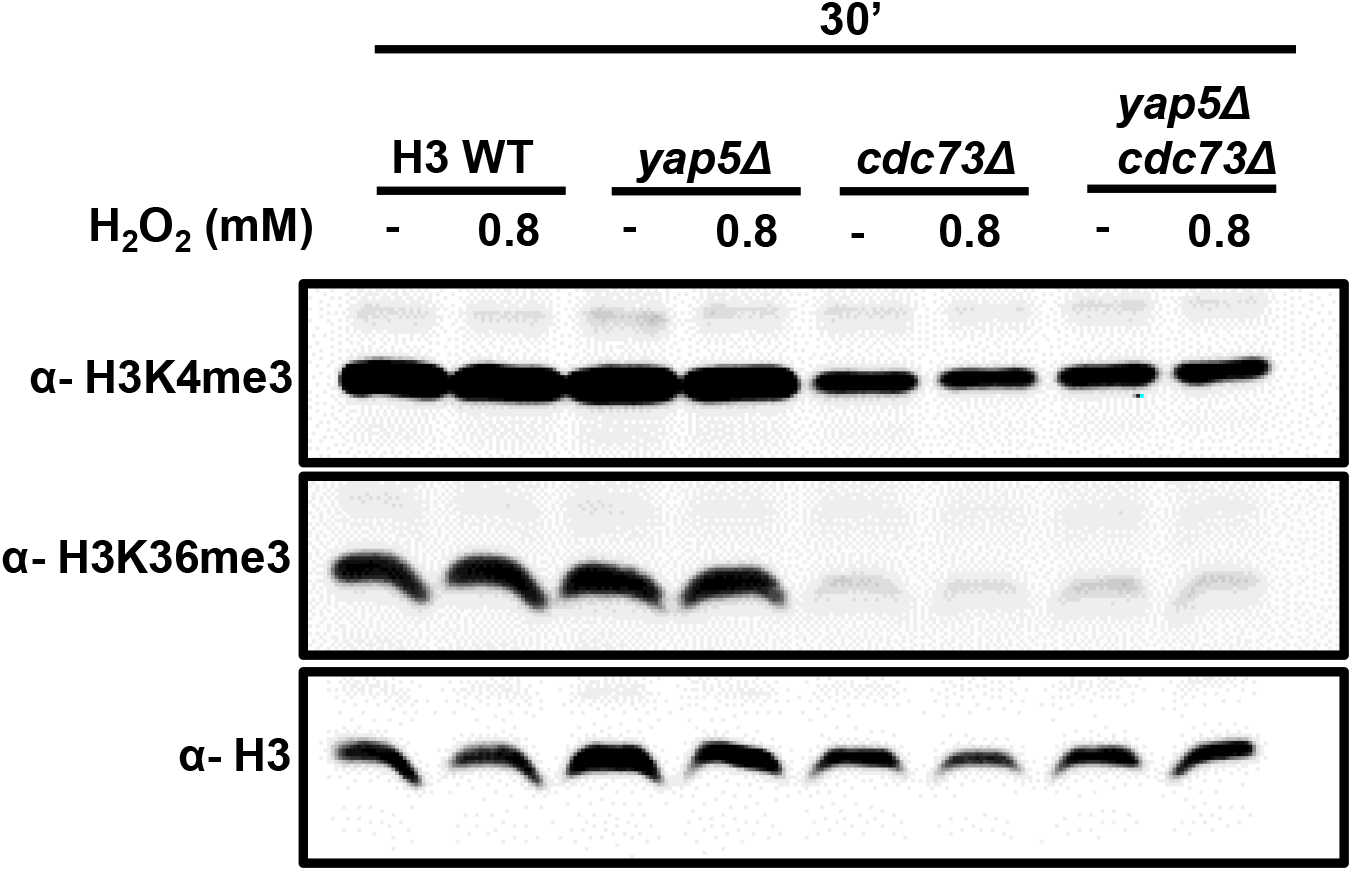
Global levels of H3K4me3 or H3K36me3 histone modifications remain unaltered under hydrogen peroxide stress. Protein extracts prepared from the H3 WT, *yap5*Δ, *cdc73*Δ or *yap5*Δ*cdc73*Δ strains untreated or H_2_O_2_ treated (0.8 mM) for 30’. Upper panel, immunoblots of α-H3K4me3 for the proteins from indicated strains. Middle panel, immunoblots of H3K36me3 for the proteins from the same strains. Lower panel, immunoblots of H3 (control) for the proteins from the same strains.

### *AIF1* gene expression levels significantly reduce in H3K36A but not in H3K4A or H3K79A mutants

Previous studies have established that Cdc73, a critical subunit of Paf1C, is important for its recruitment to the chromatin. The complex helps in the transcription of the genes by aiding H3K4 or K79 methylation [13, 15]. Additionally, the Cdc73 promotes H3K36me3 via a methyltransferase, Set2p [39]. Thus, we argued that substituting the lysine residues at these locations may impact the *AIF1* gene expression level. To this end, we grew the H3WT, H3K4A, H3K36A or H3K79A (Fig. S4 shows reconfirmation of the mutant strains through western blot) mutants till the exponential phase and either kept them untreated or treated them with 0.8 mM H_2_O_2_ for 30’. We harvested the cells, extracted RNA and then checked the levels of the *AIF1* gene relative to the *ACT1* control. As shown in Fig. 5A, B, compared to the H3WT (1.00±0.00 and 19.28±5.56 for UT and 0.8 mM H_2_O_2_, respectively), the H4K4A (1.03±0.11 and 16.65±2.58 for UT and 0.8 mM H_2_O_2_, respectively) showed almost no change in the expression pattern of the gene. The H3K79A showed subtle but significant induction of the gene compared to the H3WT in UT condition, indicating that the residue might be essential for keeping the gene level under control in the absence of stress (1.34±0.01 and 20.34±1.73 for UT and 0.8 mM H_2_O_2_, respectively). The mutant H3K36A expression analysis showed a drastic and significant reduction in the gene level in UT (0.13±0.08) or 0.8 mM H_2_O_2_ treatment (2.16±0.45). Thus, the results indicate that the H3K36 residue is critical in mediating the *AIF1* gene expression level.

**Figure 5.**
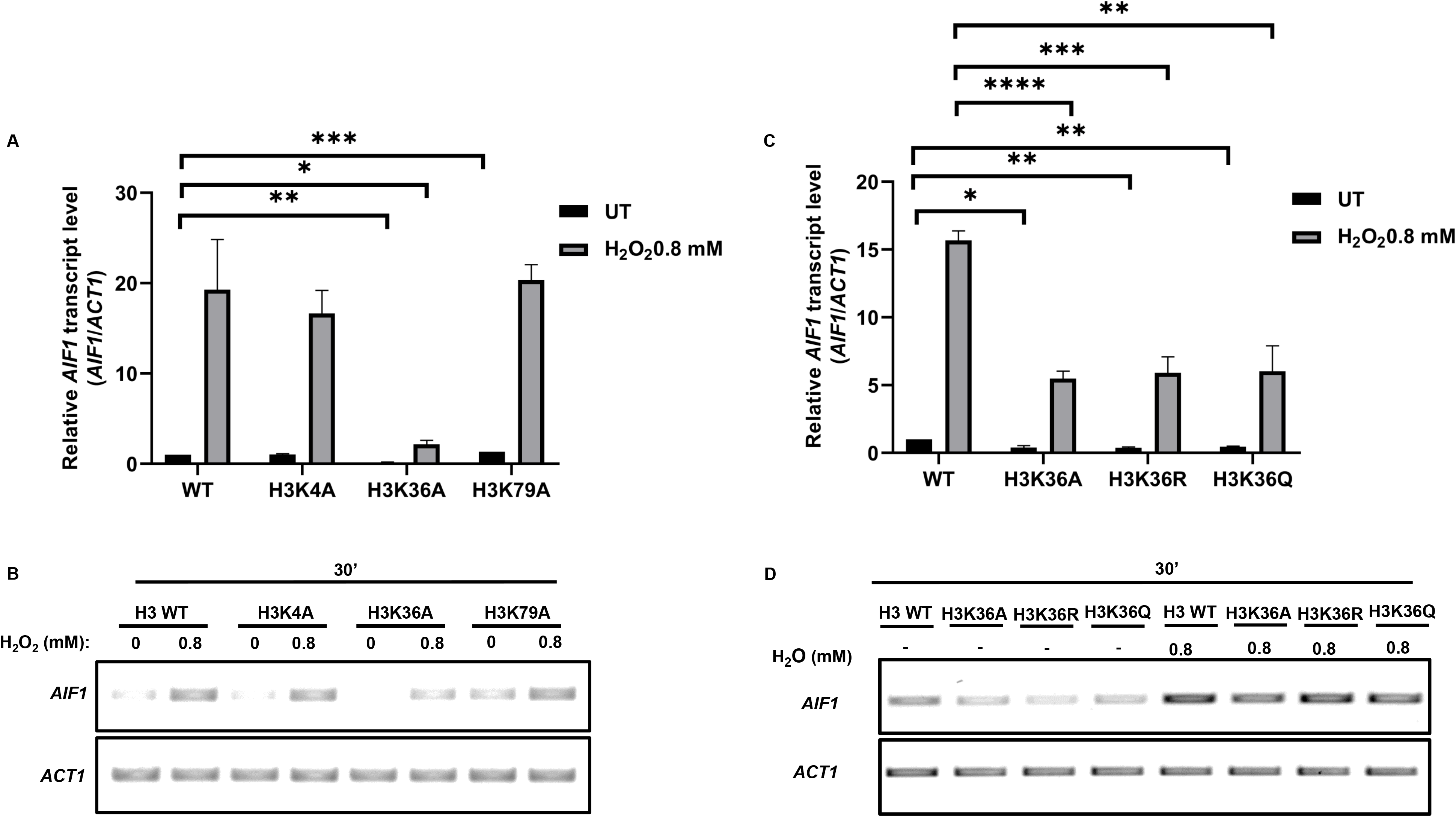
*AIF1* gene expression levels significantly reduce in H3K36A but not in H3K4A or H3K79A mutants. **A,** *AIF1* mRNA level was examined through real-time-PCR (RT PCR) in the wild-type (H3 WT), and mutants (K3K4A, H3K36A and H3K79A) under untreated (UT) or H_2_O_2_ treated (0.8 mM) for 30’, the relative expression with respect to *ACT1* (control) gene is shown. **B,** Same as A, but the results show semiquantitative data for better visualisation. **C,** *AIF1* mRNA level was examined through real-time-PCR (RT PCR) in the wild-type (H3 WT) and mutants (K3K36A, H3K36R and H3K36Q) under untreated (UT) or H_2_O_2_ treated (0.8 mM) for 30’. The relative expression with respect to the *ACT1* (control) gene is shown. **D,** Same as C, but the results show the semiquantitative data for better visualisation. Statistical analysis was performed using the Student’s t-test (in GraphPad Prism 8 GraphPad Software, Inc.). Values shown are the mean ± SD of three biological replicates (n = 3). ∗∗∗p < 0.001, ∗∗p (0.001 to 0.01), ∗p (0.01 to 0.05)

### The mutants of H3K36 residue viz. H3K36A, H3K36R and H3K36Q show significantly reduced levels of the *AIF1* gene

We next tried to decipher the impact of different mutant forms of H3K36 viz. H3K36A, H3K36R and H3K36Q on the *AIF1* gene expression levels. To this end, we grew H3WT and the H3K36 mutants (Fig. S4 shows reconfirmation of the mutant strains through western blot) till the exponential phase and either left them untreated or treated them with 0.8 mM H_2_O_2_ for 30’. The cells were harvested, RNA was extracted, and the gene expression was checked with the cDNA prepared from the samples. We found that all the variants of H3K36 exhibit a drastic and significant decrease in both the basal and hydrogen peroxide-induced levels of the *AIF1* gene (Fig. 5C, D). Compared to the H3WT, *AIF1* gene level for UT (1.00±0.00) or 0.8 mM H_2_O_2_ (15.67±0.70), the H3K36A showed (0.38±0.15 and 5.48±0.56 for UT and 0.8 mM H_2_O_2_, respectively), the H3K36R showed (0.37±0.07 and 5.91±1.18 for UT and 0.8 mM H_2_O_2_, respectively), and the H3K36Q showed (0.45±0.05 and 6.02±1.89 for UT and 0.8 mM H_2_O_2_, respectively). Thus, the results highlight the significance of the H3K36 residue and, therefore, the H3K36me3, which may play a crucial role in regulating the *AIF1* gene level.

## Discussion

Yeast cells exhibit diverse responses to the hydrogen peroxide stress [40]. Here, we show the regulation of a critical apoptotic gene, *AIF1*, for the first time. First, we established that the earliest response (15’) to the hydrogen peroxide selectively upregulates the *AIF1* gene expression level but not others. However, at 45’, the *BXI1* expression is significantly induced at 0.8 mM H_2_O_2_. This suggests that the role of different apoptotic genes becomes crucial during prolonged stress. Thus, it would be interesting to study this aspect further, taking both early and later time points. Since hydrogen peroxide stress significantly increases the reactive oxygen species (ROS) levels [40], studying the ROS levels in the different deletion backgrounds would give insights into the correlation between *AIF1* gene and ROS generation within the cells. Studies show that the Aif1p degrades the DNA during yeast apoptosis [5]. Thus, the effect of Aif1 regulator deletions on DNA fragmentation during apoptosis can be studied in detail to understand the apoptotic response by these cells (Fig. 6). Earlier, we demonstrated that acetic acid treatment in H3WT yeast cells but does not significantly induce the *AIF1* gene expression level [41]. Thus, the effect of hydrogen peroxide on the induction of *AIF1* gene expression is unique.

**Figure 6.**
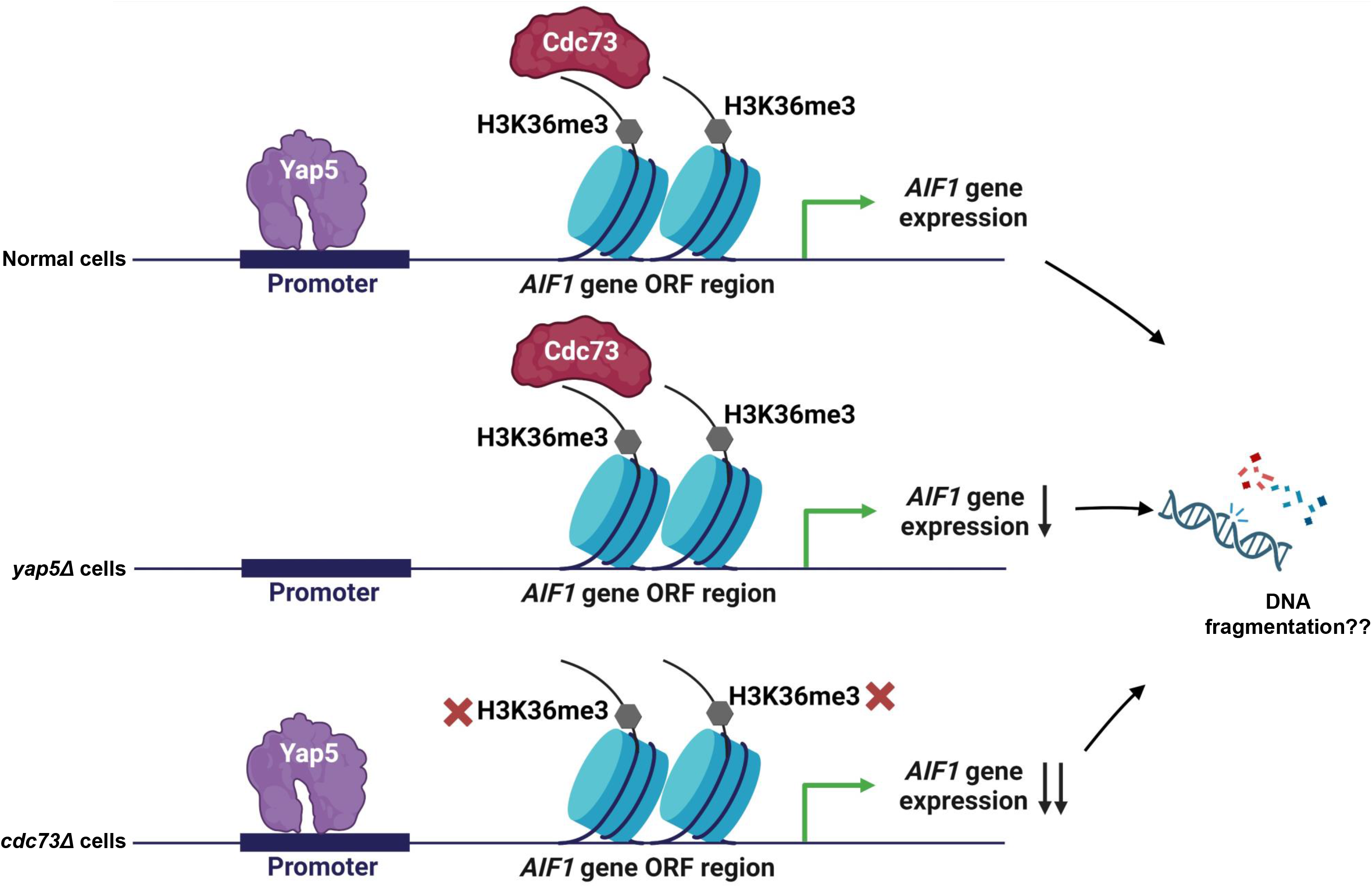
*AIF1* gene expression regulation in *Saccharomyces cerevisiae*. The wild-type cells exhibit induced levels of *AIF1* gene expression upon hydrogen peroxide stress. However, when Yap5 is deleted, induction of the gene expression is reduced. Deletion of Cdc73 majorly reduces the *AIF1* gene expression under stress, which it probably does by preventing the H3K36me3 within the ORF of the *AIF1* gene, as has been reported for other genes regulated by the Cdc73. Additionally, since Aif1p has been shown to participate in DNA fragmentation during apoptotic treatment by hydrogen peroxide, further studies are required for in-depth analyses of the DNA stability in the absence of these two regulatory factors.

Cdc73 is part of the Paf1C and significantly maintains the H3K4me3 and H3K36me3 levels in *Saccharomyces cerevisiae* [16]. We observed that the deletion of Cdc73 or mutation in H3K36 residue, either in the form of substitution with A, R or Q, severely compromises the basal and H_2_O_2_-induced gene expression levels. Studies show that Cdc73 deletion reduces the Paf1C occupancy on the transcribed genes. Also, the deletion of Rtf1, a component of Paf1C, reduces the H3K36me3 levels, which further reduces with the *cdc73*Δ. Thus, Rtf1 and Cdc73 promote the H3K36me3 together. In our study, the *cdc73*Δ does not wholly abrogate the induction of the *AIF1* gene expression, which leaves an open question about the role of Rtf1p in the *AIF1* regulation plausibly via the H3K36me3 maintenance. Further, exhaustive CHIP-based experiments on the *AIF1* transcribed regions will provide a detailed picture of the *AIF1* gene regulation.

Unexpectedly, the H3K79A, which essentially abrogates the methylation at K79 residue, increases the basal *AIF1* gene expression level. Further investigation will help understand the reason behind the observation. Studies have shown that Yap5 regulates the *AIF1* gene expression level during heat stress [37]. In this study, we, for the first time, have established the role of Yap5 in regulating the *AIF1* gene expression during H_2_O_2_-mediated stress. CHIP experiments on the Yap5 binding of the *AIF1* promoter region under hydrogen peroxide stress would provide a detailed mechanism of its gene regulation.

Thus, the revelation of hydrogen peroxide-induced *AIF1* gene expression and its regulation being dependent on H3K36me3 can pave the way for understanding apoptotic gene regulation through the lens of histone modifications.

## Material and Methods

### Culture conditions and chemical stocks used for treatment

Yeast strains (detailed in Table S1) were optimally cultured at 30 °C and 200 rpm shaking conditions maintained in the Brunswick incubator shaker. The strains were regularly maintained in synthetic complete (SC) media (0.18% amino acid mix, 0.17% yeast nitrogen base, 0.5% ammonium sulphate, 2% glucose and 2% agar for plates). For studying the response of hydrogen peroxide, a library created by the Boeke group was utilised [42]. The experiments with acetic acid were performed with YPD (pH maintained at 4.5 by adding HCl) before autoclaving [43]. Treatment with acetic acid was carried out using a stock of concentrated acetic acid (8.7 M, pH 4.5) made using glacial acetic acid and filter-sterilized. Hydrogen peroxide treatment was carried out using a fresh 1M stock of hydrogen peroxide made in sterile Milli-Q (MQ).

### Growth assays

Yeast strains were tested in various conditions using either drop-test assays or growth curves, as detailed below:

### Drop-test assays

Overnight-grown yeast cultures (primary culture) were harvested and used for drop-test assays. Briefly, an equal number of cells (with an absorbance of 1 at 600 nm) were subjected to serial dilutions of 10^-1^, 10^-2^, and 10^-3^. These dilutions were gently vortexed to ensure uniform mixing, and from each dilution, 2.5 μL (for acetic acid) or 3 μL (for others) of culture were individually spotted on plates. The spots were incubated after being air-dried at a temperature of 30 °C.

### Growth Curve Analysis

Primary cultures were normalised to an OD of 0.2 at 600 nm using media and were left untreated or treated with hydrogen peroxide or acetic acid and seeded into 96 well plates (Tarsons). These plates were incubated in the plate reader set to measure OD at 600 nm at an interval of 30 min.

### Chronological life span (CLS) assay

Chronological life span was performed using the previously described method [44] with some modifications. Saturated cultures of H3 WT and various deletions were serially diluted, and aliquots with an equal number of cells (1000) were plated on SC agar and incubated for 4 days at 30 °C. Post the incubation period, colonies were counted, and cellular viability was determined. From the same culture dilutions, drop-test assays were performed parallelly.

### Protein extraction (TCA precipitation method)

Whole-cell extracts from yeast cells were prepared following the TCA method [45] with few alterations. Briefly, the yeast cells harvested under untreated/treated conditions were washed with 20% TCA solution (made in sterile MQ) and kept at −80 °C until further use. For extraction, cell pellets were resuspended in 20% TCA, and 200 μL glass beads (1mm) were added, after which the cells were lysed using a multi-vortex mixer. The precipitated protein was pelleted by centrifugation at 6000 rpm and washed with 0.5 M Tris–Cl (pH 7.5). Finally, the pellet was boiled in 1× loading buffer, and the protein supernatant was used for the western blot.

### Western blotting

After the proteins were resolved using SDS-PAGE, they were transferred to the nitrocellulose membrane using the transfer buffer (Tris–Cl (pH 7.5), glycine, methanol and SDS) for 90 min at 4 °C. The immunoblot containing the proteins was blocked in 2.5% bovine serum albumin (made in 1X TBST) (BSA). The immunoblot was incubated with primary antibody at room temperature for 90 min, following which it was washed with 1X TBST to remove non-specific binding. The immunoblot was then probed with a secondary antibody for 45 min at room temperature and washed using 1X TBST. Finally, the dried immunoblot was visualised using the LI-COR infrared imaging system. Primary and secondary antibodies used for immunoblotting were as follows: α-H3 polyclonal antiserum from rabbit (Sigma, H0164), α-H3K4Me3 (Cell Signaling), α-H3K36Me3 and goat anti-rabbit IgG secondary antibody (Invitrogen, catalogue no.: A32734;).

### DNA isolation from yeast

As described earlier, cells were harvested from overnight-grown cultures for DNA extraction [46]. Briefly, the harvested cells were mixed with 0.2% SDS (300 μL), vortexed, and boiled for 15 min at 100 LC. Next, NaCl (100 mM) was added to the mixture, following which the contents were vortexed and centrifuged at 10,000 rpm for 10 min. The supernatant was transferred to a new vial. The lysate was mixed with an equal volume of phenol (pH 8): chloroform: isoamyl alcohol (25:24:1) and centrifuged for 10 min at 13,000 rpm. The aqueous phase was added to a fresh tube with absolute ethanol for precipitation. DNA obtained post centrifugation was air dried and resuspended in MQ and transferred to 4 LC. The manufacturer’s instructions (KAPA Taq DNA Polymerase; catalogue no.: KK1015) were followed for the semiquantitative PCR to check the absence or presence of different genes (Table S2).

### RNA extraction and quantitative real-time PCR of genes

The untreated/treated cultures were harvested after the cells were grown till the exponential phase, following which RNA was extracted using the hot phenol method [47]. RNA samples were normalised to the same concentration of 250 ng/μL and used to synthesise the complementary DNA using an iScript cDNA synthesis kit (Bio-Rad, catalogue no.: 1708891) following the manufacturer’s instructions. RT–PCRs of different genes using respective primers (Table S2) were performed using ABI-7300 RT–PCR with Sequence Detection System v1.4 (Applied Biosystems). Relative mRNA expression was calculated using the previously established 2−ΔΔCT method [48]. The relative expression fold changes were calculated using the *ACT1* gene as a control. Data are represented as mean ± SD of three replicates.

### Transformation and gene knockout

#### 1 Competent cell preparation

Overnight YPD-grown cultures at 30°C were diluted to a 0.2 OD in 30 mL YPD the next day and grown to 0.8 – 1 OD. The cells were harvested and washed with distilled water and lithium acetate buffer at this stage. Following the washes, the cell pellet was resuspended in lithium acetate buffer and incubated for 30 min at room temperature.

#### 2 Gene knockout

Templates for knocking out the different genes were generated through a 2-step PCR of pRS405 or pRS404 plasmid and the primers specific for *AIF1* KO [49]. The purified product was used as a template for transforming yeast cells [50], and the positive colonies were selected using SC-Leu plates. The knockout was confirmed through PCR amplification of the gene using corresponding primers (Table S2).

## Data availability

All the relevant data are contained within the manuscript and the supporting file.

## Supporting Information

This article contains supporting information.

## Acknowledgements

We acknowledge all lab members for their suggestions.

## Author contributions

N.S. and R.S.T. conceptualisation; N.S. and S.A. investigation, N.S. methodology, validation, and formal analysis; R.S.T supervision; N.S. writing-original draft; N.S. and R.S.T. writing – review & editing; R.S.T. project administration; N.S., S.A. and R.S.T. funding acquisition.

## Funding and additional information

This work was supported by funds from the Council of Scientific and Industrial Research, Government of India (CSIR Grant No. 38(1438)/18/EMR-II) and IISER Bhopal. N.S. and S.A. received fellowship from IISER Bhopal.

## Conflict of interest

The authors declare that they have no conflicts of interest with the contents of this article.

## Abbreviations

(*AIF1*): Apoptosis Inducing Factor
(Yap5">): Yeast AP-1
(Cdc73): cell division cycle
(H3): histone 3
(H3K36me3): H3K36 trimethylation
(H3K4me3): H3K4 trimethylation

## Notes

### Competing Interest Statement

The authors have declared no competing interest.

## References

1. Fahrenkrog B, Sauder U, Aebi U (2004). The S. cerevisiae HtrA-like protein Nma111p is a nuclear serine protease that mediates yeast apoptosis. J Cell Sci 117(Pt 1): 115–126. doi: 10.1242/jcs.00848

2. Madeo F, Herker E, Maldener C, Wissing S, Lachelt S, Herlan M, Fehr M, Lauber K, Sigrist SJ, Wesselborg S, Frohlich KU (2002). A caspase-related protease regulates apoptosis in yeast. Mol Cell 9(4): 911–917. doi: 10.1016/s1097-2765(02)00501-4

3. Buttner S, Eisenberg T, Carmona-Gutierrez D, Ruli D, Knauer H, Ruckenstuhl C, Sigrist C, Wissing S, Kollroser M, Frohlich KU, et al (2007). Endonuclease G regulates budding yeast life and death. Mol Cell 25(2): 233–246. doi: 10.1016/j.molcel.2006.12.021

4. Cebulski J, Malouin J, Pinches N, Cascio V, Austriaco N (2011). Yeast Bax inhibitor, Bxi1p, is an ER-localized protein that links the unfolded protein response and programmed cell death in Saccharomyces cerevisiae. PLoS One 6(6): e20882. doi: 10.1371/journal.pone.0020882

5. Wissing S, Ludovico P, Herker E, Buttner S, Engelhardt SM, Decker T, Link A, Proksch A, Rodrigues F, Corte-Real M, et al (2004). An AIF orthologue regulates apoptosis in yeast. J Cell Biol 166(7): 969–974. doi: 10.1083/jcb.200404138

6. Pimentel C, Vicente C, Menezes RA, Caetano S, Carreto L, Rodrigues-Pousada C (2012). The role of the Yap5 transcription factor in remodeling gene expression in response to Fe bioavailability. PLoS One 7(5): e37434. doi: 10.1371/journal.pone.0037434

7. Mollapour M, Fong D, Balakrishnan K, Harris N, Thompson S, Schuller C, Kuchler K, Piper PW (2004). Screening the yeast deletant mutant collection for hypersensitivity and hyper-resistance to sorbate, a weak organic acid food preservative. Yeast 21(11): 927–946. doi: 10.1002/yea.1141

8. Shi X, Finkelstein A, Wolf AJ, Wade PA, Burton ZF, Jaehning JA (1996). Paf1p, an RNA polymerase II-associated factor in Saccharomyces cerevisiae, may have both positive and negative roles in transcription. Mol Cell Biol 16(2): 669–676. doi: 10.1128/MCB.16.2.669

9. Wade PA, Werel W, Fentzke RC, Thompson NE, Leykam JF, Burgess RR, Jaehning JA, Burton ZF (1996). A novel collection of accessory factors associated with yeast RNA polymerase II. Protein Expr Purif 8(1): 85–90. doi: 10.1006/prep.1996.0077

10. Rondon AG, Gallardo M, Garcia-Rubio M, Aguilera A (2004). Molecular evidence indicating that the yeast PAF complex is required for transcription elongation. EMBO Rep 5(1): 47–53. doi: 10.1038/sj.embor.7400045

11. Tous C, Rondon AG, Garcia-Rubio M, Gonzalez-Aguilera C, Luna R, Aguilera A (2011). A novel assay identifies transcript elongation roles for the Nup84 complex and RNA processing factors. EMBO J 30(10): 1953–1964. doi: 10.1038/emboj.2011.109

12. Mueller CL, Porter SE, Hoffman MG, Jaehning JA (2004). The Paf1 complex has functions independent of actively transcribing RNA polymerase II. Mol Cell 14(4): 447–456. doi: 10.1016/s1097-2765(04)00257-6

13. Krogan NJ, Dover J, Wood A, Schneider J, Heidt J, Boateng MA, Dean K, Ryan OW, Golshani A, Johnston M, et al (2003). The Paf1 complex is required for histone H3 methylation by COMPASS and Dot1p: linking transcriptional elongation to histone methylation. Mol Cell 11(3): 721–729. doi: 10.1016/s1097-2765(03)00091-1

14. Ng HH, Dole S, Struhl K (2003). The Rtf1 component of the Paf1 transcriptional elongation complex is required for ubiquitination of histone H2B. J Biol Chem 278(36): 33625–33628. doi: 10.1074/jbc.C300270200

15. Ng HH, Robert F, Young RA, Struhl K (2003). Targeted recruitment of Set1 histone methylase by elongating Pol II provides a localized mark and memory of recent transcriptional activity. Mol Cell 11(3): 709–719. doi: 10.1016/s1097-2765(03)00092-3

16. Amrich CG, Davis CP, Rogal WP, Shirra MK, Heroux A, Gardner RG, Arndt KM, VanDemark AP (2012). Cdc73 subunit of Paf1 complex contains C-terminal Ras-like domain that promotes association of Paf1 complex with chromatin. J Biol Chem 287(14): 10863–10875. doi: 10.1074/jbc.M111.325647

17. Heintzman ND, Stuart RK, Hon G, Fu Y, Ching CW, Hawkins RD, Barrera LO, Van Calcar S, Qu C, Ching KA, et al (2007). Distinct and predictive chromatin signatures of transcriptional promoters and enhancers in the human genome. Nat Genet 39(3): 311–318. doi: 10.1038/ng1966

18. Ruthenburg AJ, Allis CD, Wysocka J (2007). Methylation of lysine 4 on histone H3: intricacy of writing and reading a single epigenetic mark. Mol Cell 25(1): 15–30. doi: 10.1016/j.molcel.2006.12.014

19. Bernstein BE, Kamal M, Lindblad-Toh K, Bekiranov S, Bailey DK, Huebert DJ, McMahon S, Karlsson EK, Kulbokas EJ, 3rd, Gingeras TR, et al (2005). Genomic maps and comparative analysis of histone modifications in human and mouse. Cell 120(2): 169–181. doi: 10.1016/j.cell.2005.01.001

20. Heintzman ND, Ren B (2009). Finding distal regulatory elements in the human genome. Curr Opin Genet Dev 19(6): 541–549. doi: 10.1016/j.gde.2009.09.006

21. Separovich RJ, Wilkins MR (2021). Ready, SET, Go: Post-translational regulation of the histone lysine methylation network in budding yeast. J Biol Chem 297(2): 100939. doi: 10.1016/j.jbc.2021.100939

22. Walter D, Matter A, Fahrenkrog B (2014). Loss of histone H3 methylation at lysine 4 triggers apoptosis in Saccharomyces cerevisiae. PLoS Genet 10(1): e1004095. doi: 10.1371/journal.pgen.1004095

23. Faucher D, Wellinger RJ (2010). Methylated H3K4, a transcription-associated histone modification, is involved in the DNA damage response pathway. PLoS Genet 6(8): doi: 10.1371/journal.pgen.1001082

24. South PF, Harmeyer KM, Serratore ND, Briggs SD (2013). H3K4 methyltransferase Set1 is involved in maintenance of ergosterol homeostasis and resistance to Brefeldin A. Proc Natl Acad Sci U S A 110(11): E1016–1025. doi: 10.1073/pnas.1215768110

25. Pokholok DK, Harbison CT, Levine S, Cole M, Hannett NM, Lee TI, Bell GW, Walker K, Rolfe PA, Herbolsheimer E, et al (2005). Genome-wide map of nucleosome acetylation and methylation in yeast. Cell 122(4): 517–527. doi: 10.1016/j.cell.2005.06.026

26. Bell O, Wirbelauer C, Hild M, Scharf AN, Schwaiger M, MacAlpine DM, Zilbermann F, van Leeuwen F, Bell SP, Imhof A, et al (2007). Localized H3K36 methylation states define histone H4K16 acetylation during transcriptional elongation in Drosophila. EMBO J 26(24): 4974–4984. doi: 10.1038/sj.emboj.7601926

27. Rao B, Shibata Y, Strahl BD, Lieb JD (2005). Dimethylation of histone H3 at lysine 36 demarcates regulatory and nonregulatory chromatin genome-wide. Mol Cell Biol 25(21): 9447–9459. doi: 10.1128/MCB.25.21.9447-9459.2005

28. Wagner EJ, Carpenter PB (2012). Understanding the language of Lys36 methylation at histone H3. Nat Rev Mol Cell Biol 13(2): 115–126. doi: 10.1038/nrm3274

29. Jha DK, Strahl BD (2014). An RNA polymerase II-coupled function for histone H3K36 methylation in checkpoint activation and DSB repair. Nat Commun 5(3965. doi: 10.1038/ncomms4965

30. Pryde F, Jain D, Kerr A, Curley R, Mariotti FR, Vogelauer M (2009). H3 k36 methylation helps determine the timing of cdc45 association with replication origins. PLoS One 4(6): e5882. doi: 10.1371/journal.pone.0005882

31. Ng HH, Feng Q, Wang H, Erdjument-Bromage H, Tempst P, Zhang Y, Struhl K (2002). Lysine methylation within the globular domain of histone H3 by Dot1 is important for telomeric silencing and Sir protein association. Genes Dev 16(12): 1518–1527. doi: 10.1101/gad.1001502

32. 32. van Leeuwen F, Gafken PR, Gottschling DE (2002). Dot1p modulates silencing in yeast by methylation of the nucleosome core. Cell 109(6): 745–756. doi: 10.1016/s0092-8674(02)00759-6

33. Ng HH, Ciccone DN, Morshead KB, Oettinger MA, Struhl K (2003). Lysine-79 of histone H3 is hypomethylated at silenced loci in yeast and mammalian cells: a potential mechanism for position-effect variegation. Proc Natl Acad Sci U S A 100(4): 1820–1825. doi: 10.1073/pnas.0437846100

34. Norris A, Boeke JD (2010). Silent information regulator 3: the Goldilocks of the silencing complex. Genes Dev 24(2): 115–122. doi: 10.1101/gad.1865510

35. Ontoso D, Acosta I, van Leeuwen F, Freire R, San-Segundo PA (2013). Dot1-dependent histone H3K79 methylation promotes activation of the Mek1 meiotic checkpoint effector kinase by regulating the Hop1 adaptor. PLoS Genet 9(1): e1003262. doi: 10.1371/journal.pgen.1003262

36. San-Segundo PA, Roeder GS (2000). Role for the silencing protein Dot1 in meiotic checkpoint control. Mol Biol Cell 11(10): 3601–3615. doi: 10.1091/mbc.11.10.3601

37. Venters BJ, Wachi S, Mavrich TN, Andersen BE, Jena P, Sinnamon AJ, Jain P, Rolleri NS, Jiang C, Hemeryck-Walsh C, Pugh BF (2011). A comprehensive genomic binding map of gene and chromatin regulatory proteins in Saccharomyces. Mol Cell 41(4): 480–492. doi: 10.1016/j.molcel.2011.01.015

38. Hu Z, Killion PJ, Iyer VR (2007). Genetic reconstruction of a functional transcriptional regulatory network. Nat Genet 39(5): 683–687. doi: 10.1038/ng2012

39. Chu Y, Simic R, Warner MH, Arndt KM, Prelich G (2007). Regulation of histone modification and cryptic transcription by the Bur1 and Paf1 complexes. EMBO J 26(22): 4646–4656. doi: 10.1038/sj.emboj.7601887

40. Tsang CK, Liu Y, Thomas J, Zhang Y, Zheng XF (2014). Superoxide dismutase 1 acts as a nuclear transcription factor to regulate oxidative stress resistance. Nat Commun 5(3446. doi: 10.1038/ncomms4446

41. Saha N, Swagatika S, Tomar RS (2023). Investigation of the acetic acid stress response in Saccharomyces cerevisiae with mutated H3 residues. Microb Cell 10(10): 217–232. doi: 10.15698/mic2023.10.806

42. Dai J, Hyland EM, Yuan DS, Huang H, Bader JS, Boeke JD (2008). Probing nucleosome function: a highly versatile library of synthetic histone H3 and H4 mutants. Cell 134(6): 1066–1078. doi: 10.1016/j.cell.2008.07.019

43. Liu X, Zhang X, Zhang Z (2014). Point mutation of H3/H4 histones affects acetic acid tolerance in Saccharomyces cerevisiae. J Biotechnol 187(116-123. doi: 10.1016/j.jbiotec.2014.07.445

44. Herker E, Jungwirth H, Lehmann KA, Maldener C, Frohlich KU, Wissing S, Buttner S, Fehr M, Sigrist S, Madeo F (2004). Chronological aging leads to apoptosis in yeast. J Cell Biol 164(4): 501–507. doi: 10.1083/jcb.200310014

45. Keogh MC, Kim JA, Downey M, Fillingham J, Chowdhury D, Harrison JC, Onishi M, Datta N, Galicia S, Emili A, et al (2006). A phosphatase complex that dephosphorylates gammaH2AX regulates DNA damage checkpoint recovery. Nature 439(7075): 497–501. doi: 10.1038/nature04384

46. Hoffman CS, Winston F (1987). A ten-minute DNA preparation from yeast efficiently releases autonomous plasmids for transformation of Escherichia coli. Gene 57(2-3): 267–272. doi: 10.1016/0378-1119(87)90131-4

47. Schmitt ME, Brown TA, Trumpower BL (1990). A rapid and simple method for preparation of RNA from Saccharomyces cerevisiae. Nucleic Acids Res 18(10): 3091–3092. doi: 10.1093/nar/18.10.3091

48. Livak KJ, Schmittgen TD (2001). Analysis of relative gene expression data using real-time quantitative PCR and the 2(-Delta Delta C(T)) Method. Methods 25(4): 402–408. doi: 10.1006/meth.2001.1262

49. Longtine MS, McKenzie A, 3rd, Demarini DJ, Shah NG, Wach A, Brachat A, Philippsen P, Pringle JR (1998). Additional modules for versatile and economical PCR-based gene deletion and modification in Saccharomyces cerevisiae. Yeast 14(10): 953–961. doi: 10.1002/(SICI)1097-0061(199807)14:10<953::AID-YEA293>3.0.CO;2-U

50. Gietz RD, Woods RA (2002). Transformation of yeast by lithium acetate/single-stranded carrier DNA/polyethylene glycol method. Methods Enzymol 350(87-96. doi: 10.1016/s0076-6879(02)50957-5

