## Supporting Information for "Cdc73 majorly regulates apoptosis-inducing factor (*AIF1*) in *Saccharomyces cerevisiae* via the H3K36 methylation"

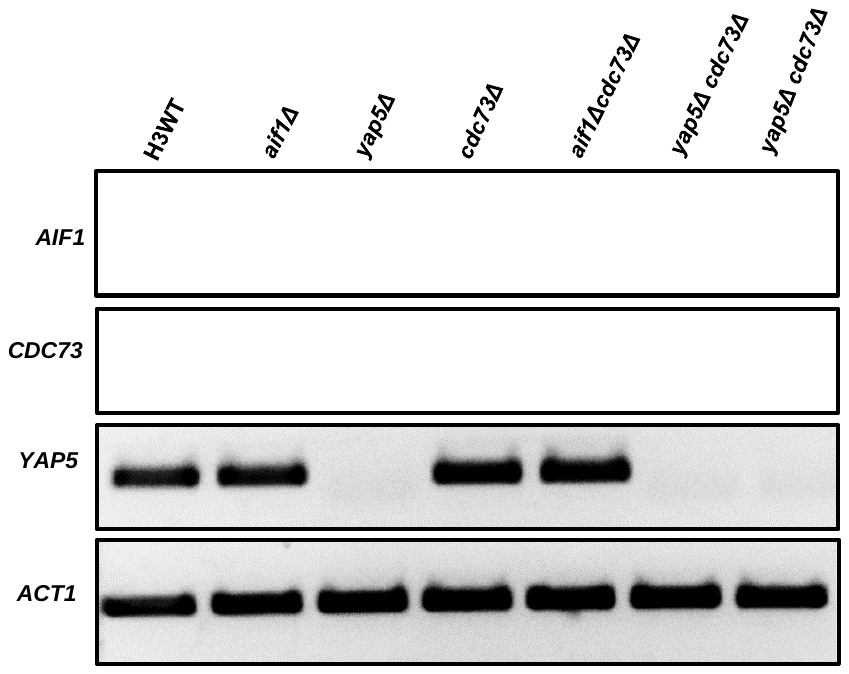


**Figure S1. Confirmation of the deletion strains.** Semi-quantitative PCR showing the *AIF1*, *CDC73*, *YAP5* and *ACT1* gene levels were examined in the DNA prepared from wild-type (H3 WT) and the indicated deletion strains. *ACT1* expression serves as the control.

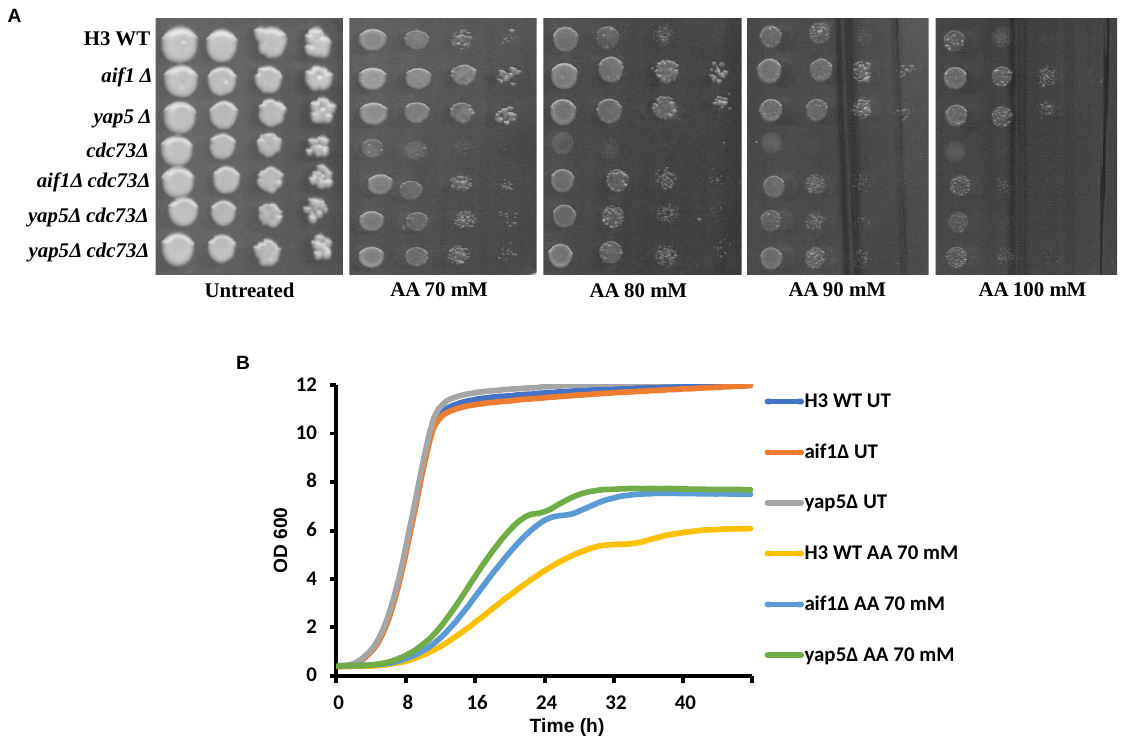

**Figure S2. Tolerance of *AIF1* gene regulators against acetic acid. A,** Spot-test assays of the wild-type (H3 WT) and gene deletion mutants in the presence of various concentrations (70, 80, 90 and 100 mM) of the acetic acid (AA) (1, 2.5, 5 nM). The spotted cells were grown for 72h and photographed. **B**, Growth curves of the indicated strains in liquid culture under untreated (UT) or acetic acid (70 mM) treated condition. The graphs reflect the absorbance values at 600 nm at every 30’ for the indicated hours in the x-axis.

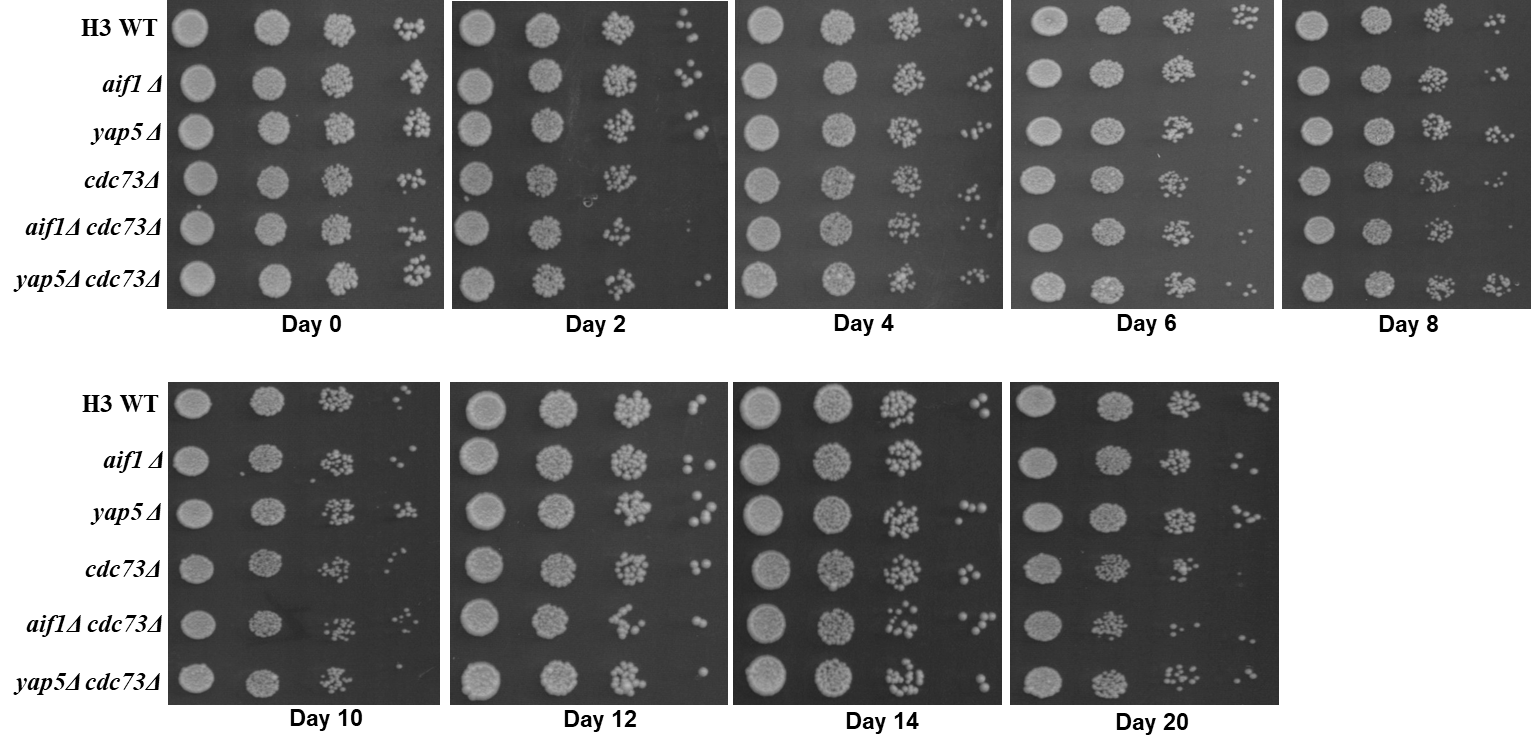

**Figure S3. Chronological life span assay with wild-type (H3 WT) and the deletion mutants.** Spot-test assay of the wildtype (H3 WT) and the deletion strains samples collected from the indicated days. The dried spots were incubated for 72h and photographed.

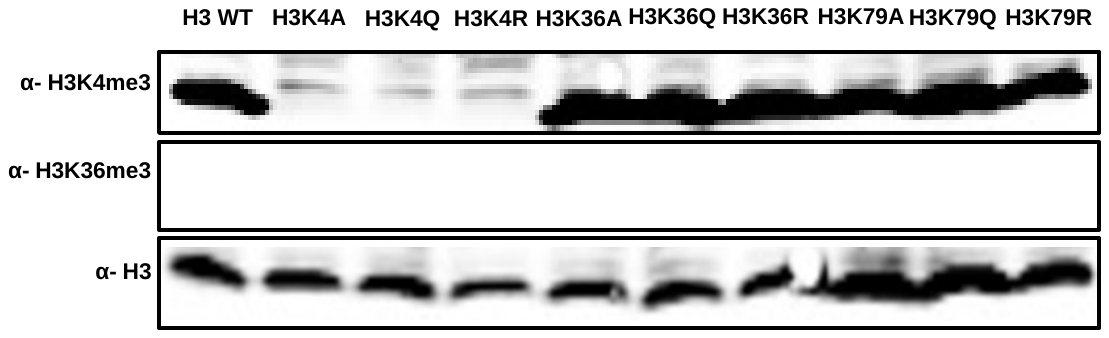

**Figure S4. Validation of H3K4A/Q/R and H3K36A/Q/R strains.** Protein extracts were prepared from the H3 WT and indicated mutant strains and western blot was performed using the indicated antibodies. Upper panel, immunoblots of α-H3K4me3 for the proteins from indicated strains. Middle panel, immunoblots of H3K36me3 for the proteins from the same strains. Lower panel, immunoblots of H3 (control) for the proteins from the same strains.

**Table S1: List of histone H3 mutants used in the study.**

**All strains are isogenic to *MATa his3Δ200 leu2Δ0 lys2Δ0 trp1Δ63 ura3Δ0 met15Δ0 can1::MFA1pr-HIS3 hht1-hhf1::NatMX4 hht2-hhf2::[HHTS-HHFS]*-URA3***

| S.No. | Strain name |
| --- | --- |
| 1 | H3 WT |
| 2 | H3K4A |
| 3 | H3K4R |
| 4 | H3K4Q |
| 5 | H3K36A |
| 6 | H3K36R |
| 7 | H3K36Q |
| 8 | H3K79A |
| 9 | H3K79R |
| 10 | H3K79Q |
| 11 | H3 *aif1Δ* |
| 12 | H3 *yap5Δ* |
| 13 | H3 *cdc73Δ* |
| 14 | H3 *cdc73Δyap5Δ* |
| 15 | H3 *cdc73Δaif1Δ* |

**Table S2: List of primers used in this study.**

| **Primers Name** | **Primers Sequence (5′-3′)** | **References** |
| --- | --- | --- |
| *ACT1*-F | TCGTCGGTAGACCAAGACAC | This study |
| *ACT1*-R | TTCTTCTGGGGCAACTCTCA | This study |
| *AIF1*-F | GCTGTCCGTTTGACGGTTTC | This study |
| *AIF1*-R | GATCAGCCCACTTTGAGCCA | This study |
| *YCA1*-F | TCCTCCACCTAACCAGCAGT | This study |
| *YCA1*-R | TTGTGCCTTTGCCTGTTCCT | This study |
| *NMA111*-F | AGTTTGGCTAAGGTCGGCTC | This study |
| *NMA111*-R | AACCACTTGAACCGCCAGAA | This study |
| *NUC1*-F | TGCAGAACCGCGAAGAGTTT | This study |
| *NUC1*-R | CCTCGATCATAGCCCGACCT | This study |
| *YAP5*-F | GCAAGTGCTGGAGGAAACCA | This study |
| *YAP5*-R | TGCAGGATAGGGTCCTGAGC | This study |
| *YAP5KO*-F | CACACAACATAAACAGTGTAACTAGCATATTATCACAGTCCTGTGCGGTATTTCACACCG | This study |
| *YAP5KO*-R | TTCAATGACGTATTTATAAGTATTAAGAAGTTCTCTCTCTAGATTGTACTGAGAGTGCAC | This study |
| *CDC73*-F | CGGTCAGAAAGGCGAGACAT | This study |
| *CDC73*-R | CTTAGCACCACGCAATGCAG | This study |
| *CDC73KO*-F | AAAAGAATAATAATTTGAGCAAGAAACTGGTGAAAAAATTCTGTGCGGTATTTCACACCG | This study |
| *CDC73KO*-R | ACTTTCAATGGCCGAAATACCATTCTTCCGTTTATCGTATAGATTGTACTGAGAGTGCAC | This study |
| *AIF1KO*-F | AGGAAAGAGCAGAGAAAGGAAGAAAGAAATTGCAAAATAT CTGTGCGGTATTTCACACCG | This study |
| *AIF1KO*-R | ATATATATATATATATATACGCTGCAGTTCATATTTTAGTAGATTGTACTGAGAGTGCAC | This study |
